# INTERPRETABLE MACHINE LEARNING UNVEILS KEY PREDICTORS AND DEFAULT VALUES IN AN EXPANDED DATABASE OF HUMAN IN VITRO DERMAL ABSORPTION STUDIES WITH PESTICIDES

**DOI:** 10.1101/2023.05.11.540395

**Authors:** D. Sarti, J. Wagner, F. Palma, H. Kalvan, M. Giachini, D. Lautenschalaeger, V. Lupianhez, J. Pires, M. Sales, P. Faria, L. Bertomeu, M. Le Bras

**Affiliations:** Ph.D in Statistics, Independent Researcher; Knoell Germany GmbH, Germany; Instituto ProHuma de Estudos Cientificos, Brazil; Corteva Agriscience, Brazil; Bayer CropScience, Brazil; Ouro Fino Quimica S.A, Brazil; Syngenta Protecao de Cultivos, Brazil; BASF S.A., Brazil; Knoell France SAS, France; Dr. Knoell Consult Schweiz GmbH, Switzerland

**Keywords:** Dermal absorption, *in vitro* human skin, non-dietary risk assessment, plant protection products, agrochemicals, pesticides, OECD 428, interpretative machine learning

## Abstract

The skin is the main route of exposure to plant protection products for operators, workers, residents, and bystanders. Assessing dermal absorption is key for evaluating pesticide exposure. The initial approach to risk assessment involves using default dermal absorption values or applying read-across data from experimental results from different formulations. In this way, to support non-dietary pesticide risk assessment focused but not limited to Brazil, this project evaluated 759 GLP-compliant *in vitro* human skin dermal absorption studies covering 25 formulation types and 248 active substances at multiple concentrations using interpretable machine learning techniques. Bayesian Additive Regression Trees – BART method indicated that Log Pow and molecular weight have the highest importance when predicting dermal absorption; both parameters exhibit moderate interaction uncertainty within each other and with formulation groups water-based and organic-solvent based and with tested form (concentrates or dilutions). The default values for each formulation group were determined using the upper bound of a non-parametric confidence interval for a specified quantile, with calculations conducted via bootstrapping methods; the proposed values correspond to the upper limit of the 95% confidence interval for the 95^th^ percentile: for concentrates, 10% for organic-solvent based, 4% for water-based and 3% for solid formulations. For dilutions, 42% for organic-solvent based, 37% for water-based and 39% for solid formulations. Organic-solvent based dermal absorption values from experimental data can be used as conservative surrogates for solid and water-based formulations. When no experimental data is available for higher spray dilutions of a given formulation type, a pro-rated correction is proposed to a 2 to 5-fold concentration difference, limited to the respective formulation group default value.

## 1. Introduction

Human skin forms a natural barrier of protection to the exogenous environment and to the absorption of exogenous substances, including pesticides, and it is comprised by three main layers: i) epidermis; ii) dermis; and iii) hypodermis. The deepest layer of the epidermis produces primary cells called keratinocytes, which, when they lose their nuclei in the process of migration to the stratum corneum, become flattened and cornified, thus forming a barrier to exogenous agents. Finally, a process of normal epidermal desquamation leads to the renewal of the cells and eliminates the residues that could be eventually trapped in the epidermis (WHO, 2006).

When an exogenous agent is in contact with the skin, the agent must penetrate the entire epidermis and the basement membrane to reach the capillaries in the dermis and the systemic circulation. Penetration may also occur via sweat- and sebaceous glands and hair follicles directly to the dermis (WHO, 2006).

Dermal absorption is one of the most relevant parameters to assess the risk following human exposure to pesticides^1^ as skin is usually the main potential route of human exposure during mixing, loading, application, and post application activities (OECD, 2019).

Most plant protection products are applied by spraying a water dilution of different concentrations of commercial formulations (Aggarwal *et al*., 2014). Exposure to diluted formulations is usually associated with a higher dermal absorption expressed in % of the applied dose as a larger % of the applied dose is solubilized in the water phase and therefore available for dermal absorption. It is therefore important to consider the concentration to avoid underestimating the internal dose after exposure to dilutions and/or neat products.

In a typical OECD test-guideline compliant *in vitro* human skin dermal absorption study (OECD 428), a pesticide (neat formulation/concentrate) and one or more spray dilutions containing – preferably – radiolabelled active substance is applied to the human skin for at least 6-10 hours to match a representative working day. Dermal absorption is determined through the material collected by continuous measurements for up to 24 hours after the initial skin contact (OECD, 2019).

In the European Union, when no experimental data are available on the formulation, the European Food Safety Authority – EFSA Guidance on Dermal Absorption (2017), defines the following default values to be used as an alternative:

- organic solvent-based^2^ or other formulations^3^: 25% for concentrates and 70 % for dilutions;
- water-based/dispersed^4^ or solid formulations^5^: 10% for concentrates and 50 % for dilutions.

In Brazil, according to the regulation *“Resolução da Diretoria Colegiada 294”* (RDC 294/2019) from Brazilian Health Regulatory Agency (ANVISA, 2019), studies to estimate *in vitro* or *in vivo* dermal absorption, or both, could be submitted. When no studies are available, the use of default values or read-across can be used on a scientific and technical basis. However, for the Brazilian laboratories, *in vitro* dermal absorption studies with human skin are an imminent and developing knowledge, in addition to uncertainties with regards to the process on the use of human tissue for studies supporting the registration of products to be commercialized.

The current publication presents findings from a large database of human skin *in vitro* dermal absorption studies with pesticides (ProHuma/ECPA dataset). It consists of a total of 759 studies obtained by evaluating 468 studies conducted mainly from 2014 to 2019 (ProHuma dataset) in combination with 291 studies that comprise the dataset from the European Crop Protection Association - ECPA, currently CropLife Europe (Aggarwal *et al*., 2014; Aggarwal *et al.,* 2015). This junction led to a database with a total of 14,803 observation values^6^. The data was analyzed using non-parametric advanced Bayesian techniques and presented via new interpretable machine learning tools for visualization of interactions and importance of predictors of dermal absorption.

This work was initiated by ProHuma, a consortium of several pesticide companies, aiming to provide the Brazilian health regulatory agency with consistent data to support the non-dietary risk assessment when specific studies are unavailable, allowing the possibility of establishing dermal absorption default values and read-across between pesticide formulations as an alternative. Utilizing robust existing data helps prevent unnecessary studies conduction, ensuring that scientific efforts are directed efficiently and effectively.

## 2. Objectives

To assess an extended dermal absorption database of *in vitro* human skin studies with pesticides using an appropriate statistical tool for non-parametric data and using cutting edge interpretable machine learning techniques to support the non-dietary risk assessment focused but not limited to Brazil by: i) determining potential relationships between dermal absorption and various factors such as tested form (concentrates or dilutions) and physical state (solid or liquid); ii) establishing dermal absorption default values and the percentiles of dermal absorption for different formulation groups and formulation types, with associated uncertainties covered by upper bound of confidence intervals and complemented by bootstrap methods; and iii) proposing read-across possibilities.

## 3. Methods

### 3.1. Data collection

Data collection from ProHuma dataset and subsequent analysis for this project were performed by qualified professionals from independent third-party consultancy companies.

As the study reports provided by the companies associated to ProHuma are considered confidential, Planitox, a Brazilian consultancy, under a confidentiality agreement with each company, was responsible for extracting the data from the studies to form the ProHuma dataset. Neither ProHuma nor any of its members had access to the reports, except for the studies authored by the companies themselves. The data was compiled into a spreadsheet following the pre-existing template of the ECPA dataset, using the same column headers.

The spreadsheet containing the information from ProHuma dataset was combined with the ECPA dataset by Knoell Germany GmbH, a European consultancy. Knoell performed a quality check on the dataset (see Appendix A from Supplemental Material).

ProHuma dataset includes studies covering 22 different formulation types and 190 active substances, and the combined ProHuma/ECPA dataset, 25 different formulation types with 248 active substances (Table 1). Formulation types that were not classified as organic solvent-based, solid, or water-based/dispersed, were grouped as “other types” as there was no specific characterization for these.

**Table 1:** Grouping of formulation types and codes used in this assessment.

| Grouping of formulation types | Formulation type<sup>(1)</sup> | Description for formulation type<sup>(1)</sup> | Number of studies (# of study identifiers) |  | Number of observations (individual values of replicate analysis) - Combined ProHuma/ ECPA dataset |
| --- | --- | --- | --- | --- | --- |
|  |  |  | Dataset ProHuma | Combined ProHuma/ ECPA dataset |  |
| Organic solvent-based | EC | Emulsifiable concentrate | 118 | 182 | 3559 |
|  | EW | Emulsion, oil in water | 8 | 18 | 368 |
|  | SE | Suspo-emulsion | 15 | 40 | 907 |
|  | DC | Dispersible concentrate | 2 | 2 | 40 |
|  | OF | Oil miscible liquids | 1 | 1 | 16 |
|  | OD | Oil-based suspension concentrates | 7 | 23 | 425 |
|  | ES | Emulsion for seed treatment | 2 | 3 | 53 |
|  | ME | Microemulsion | 6 | 8 | 209 |
| Water-based /dispersed | SL | Soluble concentrate | 21 | 33 | 667 |
|  | SC | Suspension concentrate | 127 | 217 | 4217 |
|  | FS | Flowable concentrate for seed treatment | 46 | 60 | 785 |
|  | CS | Capsule suspension | 13 | 14 | 174 |
|  | ZC | Mixture of capsule suspension and suspension concentrate | 4 | 7 | 129 |
|  | FL (=SC), SD | Flowable | 1 | 1 | 8 |
|  | LS | Seed treatment | 1 | 1 | 32 |
| Solids | WP | Wettable powder | 4 | 11 | 179 |
|  | WG/WDG | Water-dispersible granules | 74 | 108 | 2154 |
|  | SG | Water-soluble granules | 4 | 8 | 126 |
|  | WS | Water dispersible powder for slurry treatment | 1 | 1 | 12 |
|  | GR | Granule | 1 | 1 | 8 |
|  | DS | Powder for dry seed treatment | 0 | 1 | 5 |
| Other types | RB | Bait, ready for use | 4 | 7 | 50 |
|  | UL | Ultra-low volume | 1 | 1 | 8 |
|  | CB | Bait concentrate | 0 | 1 | 7 |
|  | GEL | - | 0 | 2 | 16 |

(1) Formulation types were available on each report and formulation descriptions and grouping were based on Manual on Development and Use of FAO and WHO Specifications for Pesticides (2016), and ABNT NBR 12679: 2018.

### 3.2. Data evaluation

Individual results for each analysis replica were used in this assessment. The data were extracted from study reports, summarized, and inserted into a Microsoft Excel (2016) spreadsheet.

The dermal absorption value was calculated in the same manner as in previous publications on this topic (EFSA, 2017; Aggarwal *et al.,* 2015):

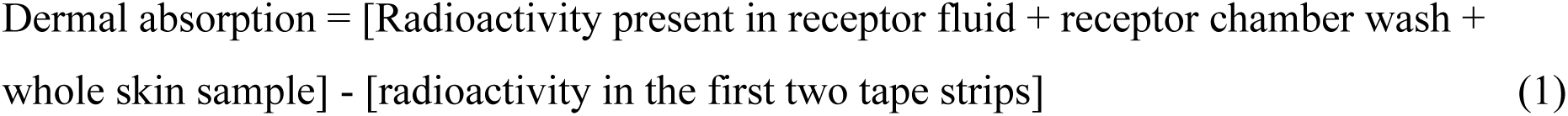

Data are presented in several types of graphs and charts, including boxplots. The different boxplots show the 25^th^ percentile (lower end of the box), the median (horizontal line within the box) and the 75^th^ percentile (upper end of the box). The 95^th^ percentile of the distributions were indicated as a triangle symbol. Additionally, Q-Q plot was used. A Q-Q plot (Quantile-Quantile plot) is a graphical tool used to compare two probability distributions by plotting their quantiles against each other. It is also known as a probability plot or normal probability plot. The quantiles are calculated in the sample data and plotted against actual values of quantiles produced by a normal distribution to check if the sample data is normal or not.

### 3.3. Strength of dataset and exclusion of studies

Both ProHuma and ECPA datasets are considered homogeneous, robust, and representative of most all plant protection products since the following criteria from the main regulatory bodies are met:

- *in vitro* human skin dermal absorption studies;
- performed with agrochemicals products;
- compliant to the OECD 428 with human skin, with the recommended recovery criteria (90-110%);
- GLP compliant;
- exposure periods of 6 to 10 hours (representative of a working day);
- period of the experiment not exceeding 24 h;
- individual values for tape strips 1 and 2 reported;
- did not have the external addition of surfactant or adjuvants.

Initially, ProHuma dataset included 488 studies; however, to meet the above criteria, 20 studies were excluded from further evaluation. Additionally, studies were excluded based on the following: formulation type was “dry residue” or “tank mixture” or conducted with a formulation that was not in its final commercial form of use. Steps involved in the data processing are presented in the Appendix A in Supplementary Material.

### 3.4. Statistical analysis and software

The spreadsheet with individual replicate analysis was read and analysed into the statistical programming language R (RStudio version 1.3.1093 (2020-09-17); R-4.0.3 (October, 2020)). The graphs, medians, and percentiles (25^th^, 75^th^ and 95^th^) were created and calculated using the RStudio and statistical functions. Graphs and boxplots were created using ggplot2.

To explore the potential relationships between dermal absorption and different factors, a machine learning method called Bayesian Additive Regression Trees – BART (Chipman *et al*., 2010) was used. BART models are particularly advantageous because they are non-parametric and require minimal assumptions about data structure having had extensive applications in economics, health care, sociology, and genetics (Sarti *et al*., 2023). BART model was fitted using dbarts R package and bartMan (Inglis *et al*., 2024), which allow to consider all variables simultaneously in a multivariate context. This approach is superior to univariate models, as it captures complex interactions and non-linear relationships among predictors. The bartMan R package (Inglis *et al*., 2024) was used to assess variable importance and interactions, providing advanced visualization tools for analysis. Specifically, Value Suppressing Uncertainty Palettes (VSUP) was utilized to create a heatmap that illustrates both the importance of variables and their interactions, incorporating uncertainty measures. These visualizations provided key insights into the factors most influencing dermal absorption and how they interact. This method allows for a comprehensive analysis, effectively identifying and quantifying the relationships between multiple predictors and the response variable (Inglis *et al*., 2024). In BART models, variable interaction captures the joint influence of two or more predictors on the response. Unlike random forests, where splits are optimized, BART employs a stochastic search, making the order of splits irrelevant. Interaction strength is identified by analysing tree structures that incorporate multiple variables. Chipman *et al*., 2010 and Kapelner and Bleich, 2016 introduced measures based on successive splitting patterns within tree-models to quantify interactions.

The quantile estimation was done using two main methods. The first method is the one proposed by EFSA, 2017, whose computational code is presented in Craig and Guillot, 2017. According to the presented code, the estimation of the quantiles is made using an R function quantile and respective upper bound by a function named ‘ub.leftCI’ that is designed to calculate the upper bound of a non-parametric confidence interval for a specified quantile.

The function operates based on the binomial distribution, which models the probability of obtaining a given quantile from the sample data. The method is particularly useful when working with empirical data that may not follow a known parametric distribution.

The second method uses bootstrapping and offers an alternative approach to estimating confidence intervals, particularly when the assumptions of the ‘ub.leftCI’ method are not met or when the sample size is inadequate. Bootstrapping can be defined as a computer-based method of random resampling technique for quantifying sampling uncertainty (Efron, 1993). Details of this function is presented in the Appendix B in Supplementary Material.

## 4. Results

The following sections will present the findings from the assessment of the extended dermal absorption database of human skin *in vitro* studies.

### 4.1 Assessment of dataset distribution

This initial assessment was performed by grouping percentage of dermal absorption results into ‘concentrates’ and ‘dilutions’. ‘Concentrates’ are the formulated products tested in their original form of commercialization, and ‘dilutions’ are the formulated products added of an appropriate diluent to reach the final use spray solution concentration as recommended on the product label.

Once the fraction of applied dose that it absorbed from finite dose experiments depends on the amount of the chemical that is initially loaded on the skin surface (Frasch *et al*., 2014), other approaches using different metrics would be possible. However, considering the currently available regulatory models for non-dietary risk assessment, the percentage of dermal absorption was the focus of this research.

A Q-Q Plot produced by the R-studio platform of the dermal absorption data from the concentrates and dilutions from the combined dataset does not follow a normal distribution (**Figure 1**). When these data are transformed to logarithmic values, the data is not normal distributed as well.

**Figure 1:**
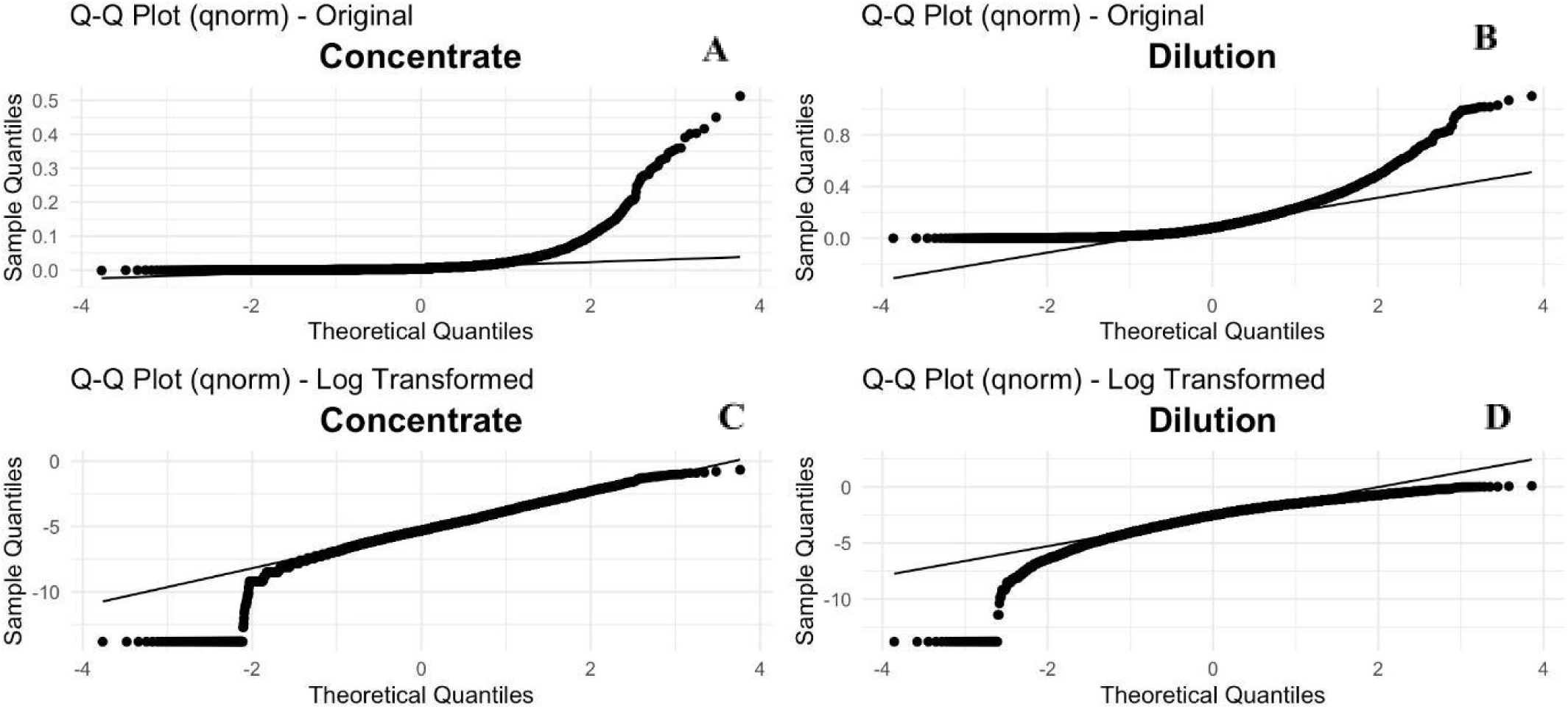
Q-Q-Plot of dermal absorption values (A, B) and Q-Q-Plot of log-transformed (C, D) for concentrates (A, C) and dilutions (B, D) of the combined ProHuma/ECPA dataset

### 4.2 Influence of different parameters on dermal absorption

The graphical analysis of variable importance (Vimp) and interaction (Vint) within the BART model (**Figure 2**) reveals critical insights into the relationships and significance of various predictors in the dataset. The following parameters were evaluated: molecular weight; log Pow; tested form (concentrates or dilutions); physical state (solid or liquid); tested concentration; grouping of formulation type (organic, water-based, solid, other types). It also highlights the associated uncertainty through the coefficient of variation (CV).

**Figure 2:**
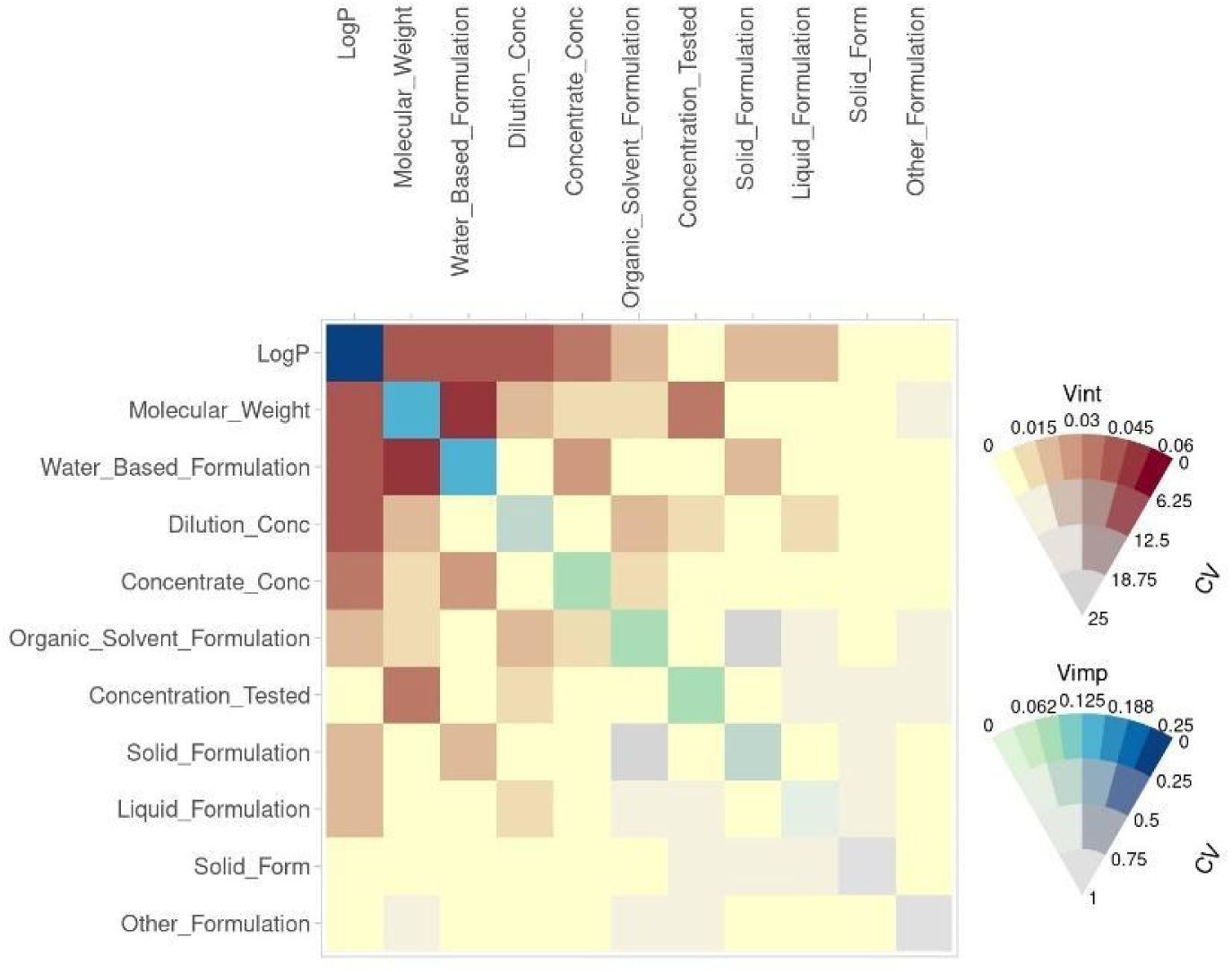
Heatmap of interaction and importance of different parameters on dermal absorption. Evaluated parameters were log Pow (*LogP*), molecular weight, formulation group (*Organic_Solvent_Formulation*; W*ater_Based_Formulation*; *Solid_Formulation*; *Other_Formulation*), tested form (*Concentrate_Conc; Dilutions_Conc*), tested concentration (*Concentration_Tested*), and physical state (*Solid_Form; Liquid_Formulation*)

The heatmap uses a combination of two metrics:

- Vint (Variable Interaction): This is shown on the right color scale using shades of red and yellow. Darker reds represent stronger interactions, while lighter shades represent weaker interactions. The uncertainty associated with each interaction is presented via the Coefficient of Variation (CV) scale in the legends; higher CVs imply higher uncertainty. It should be read and interpreted horizontally;
- Vimp (Variable Importance): Shown with shades of blue and green. Darker blues represent higher importance of the variables, while lighter shades represent lower importance. It should be read and interpreted diagonally.

The matrix itself shows pairwise relationships between parameters, with the strength of the importance and uncertainty reflected in the color intensity. The rows and columns correspond to the parameters under evaluation.

From Figure 2, it can be observed that Log Pow and molecular weight have the highest importance (deep and light blue on the diagonal) when predicting dermal absorption, which is aligned with the knowledge that these factors have a key role in cell penetration (WHO, 2006). Both parameters exhibit moderate interaction uncertainty within each other and with formulation groups (water-based and organic) and tested form (concentrates or dilutions), as indicated by the reddish shades. Concentration tested and molecular weight also presents moderate interaction uncertainty.

Parameters such as physical state (liquid or solid) and formulation groups solid and other types have lower importance and interaction uncertainty in predicting dermal absorption.

The concept of importance and interaction serves as a statistical tool that helps understand the influence of different variables on a model’s outcomes. However, it’s important to note that there isn’t always a straightforward connection between the levels of importance or interaction and the actual physicochemical properties of the substances involved. This means that while a variable might show high statistical importance in predicting a certain outcome (like dermal absorption), it does not necessarily reflect a direct causal relationship with the underlying physical or chemical properties. So, while BART is well-suited for capturing interactions non-parametrically and excels at modelling complex interactions, it does not explicitly disentangle cause-and-effect relationships between variables. Therefore, when interpreting the results, it is important to account for this limitation and acknowledge that the relationships within the variables may be complex and indirect.

Finally, BART provides robust and flexible nonlinear modeling capabilities but does not inherently offer mechanistic explanations. Unlike parametric models, BART does not yield explicit coefficients or functional forms that describe the relationships between predictors and outcomes unless complemented by interpretable machine learning techniques (Chipman *et al*., 2010; Kapelner and Bleich, 2016, Inglis *et al*., 2024). This constraint limits its ability to provide direct interpretability and mechanistic insights, which may be necessary for certain applications.

Considering that the main objective of this work is to support non-dietary risk assessment in Brazil, it was decided to use the following categorization of pesticides to calculate the 95^th^ percentile to be harmonized with EFSA, 2017, an international well-known dermal absorption guideline: formulations were classified as organic-solvent based, water-based and solids (any other formulation that doesn’t fit into these three categories were classified as ‘other types’), and separated into other 2 groups: concentrates and dilutions (as defined in 4.1.). This approach was adopted with the intention of contributing to a common regulatory terminology across regions.

### 4.3. Effect of formulation type

#### 4.3.1. Concentrates

The potential impact that individual formulation type and formulation-type groupings have on dermal absorption from concentrates was assessed (**Figure 3**).

**Figure 3:**
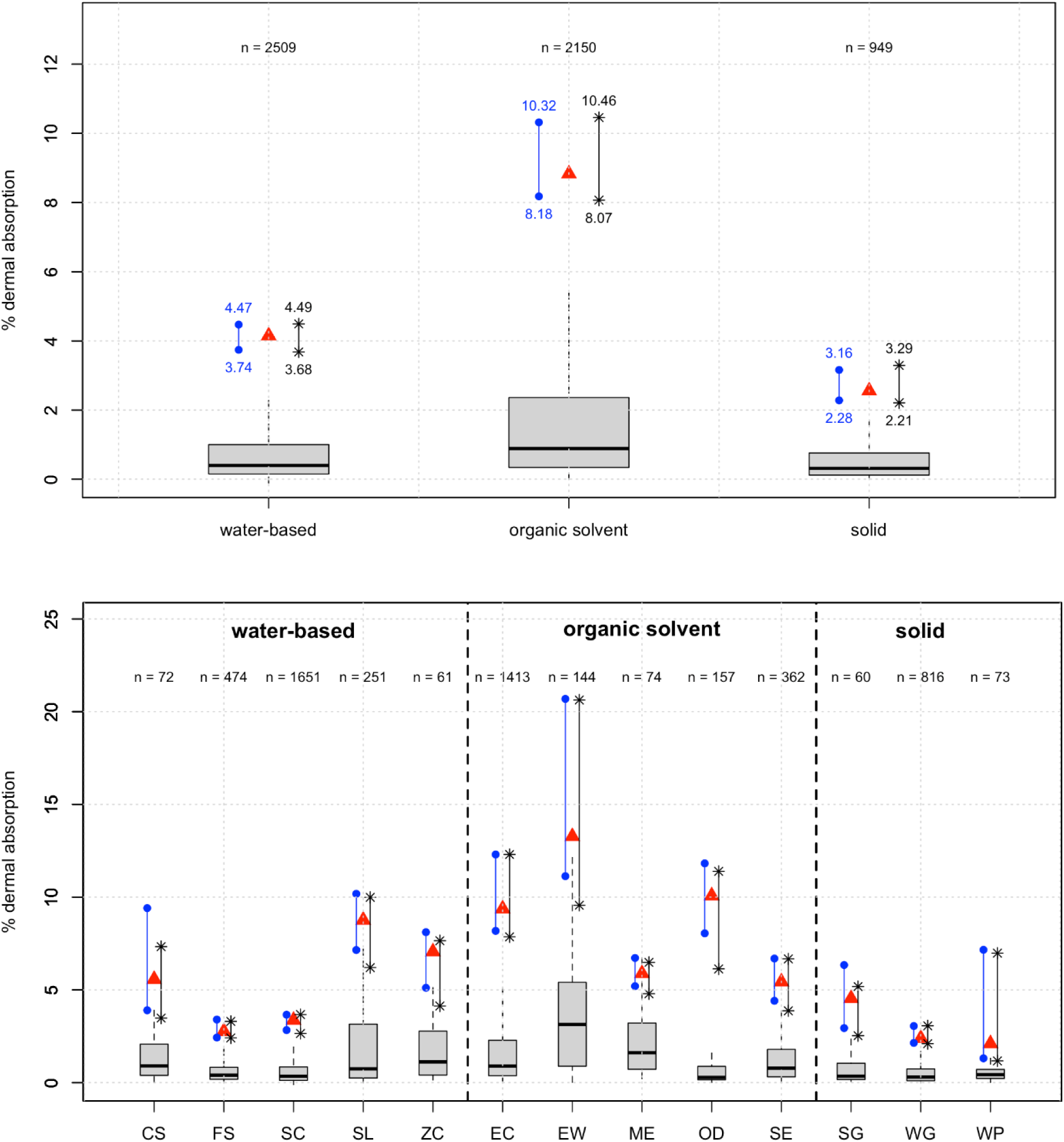
Dermal absorption of concentrates by grouped formulations (top) and formulation type (bottom) with superimposed 95^th^ percentiles (red symbol, Craig and Guillot, 2017 method) and 95% Confidence Interval of 95^th^ percentile (blue dots: Craig and Guillot, 2017 method; black asterisk: Bootstrap method). Only formulation types with minimum of 7 studies were included for this evaluation.

To compare differences of the 95^th^ percentile between formulation groups and within each group, Bootstrap method was used to calculate lower and upper 95% confidence interval of the 95^th^ percentile (**Figure 3**). In this approach, if the confidence intervals overlap significantly, the 95^th^ percentiles may not differ significantly between groups or formulation types under assessment. Appendix C in Supplementary Material presents the 95^th^ percentile (95% upper confidence interval) using Bootstrap for each formulation type.

For concentrates, the lack of overlap between the 95% confidence intervals of the 95^th^ percentiles indicates that these values are significantly different from each other. Within each formulation group, EW shows the highest value among organic solvent formulations, WP among solid formulations, and SL among water-based formulations.

#### 4.3.2. Dilutions

As presented for the concentrates, the potential impact that an individual formulation type and formulation-type groupings have on dermal absorption from spray dilutions was also assessed (**Figure 4**).

**Figure 4:**
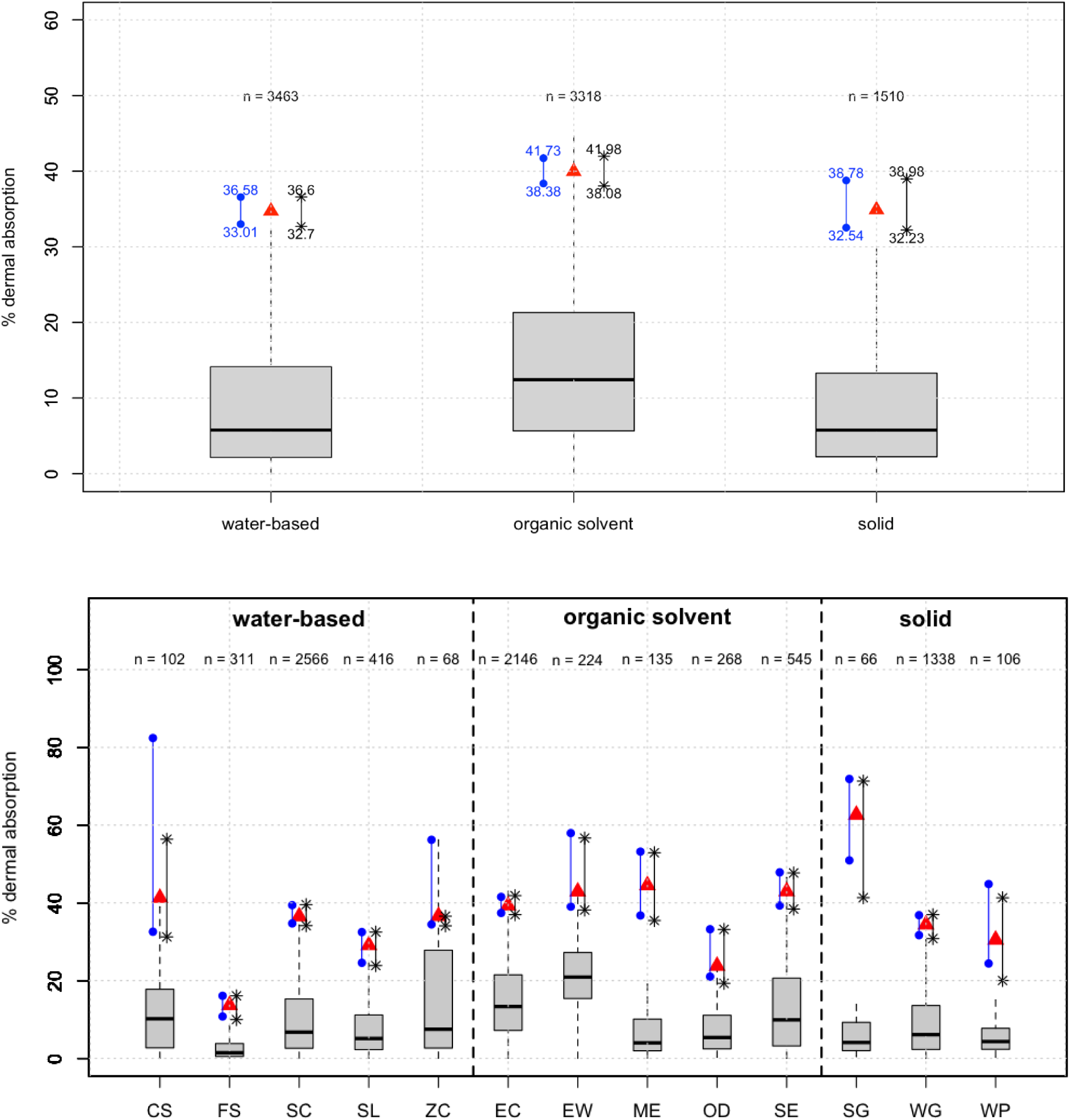
Dermal absorption of dilutions by grouped formulation (top) and formulation type (bottom) with superimposed 95^th^ percentiles (red symbol, Craig and Guillot, 2017 method) and 95% Confidence Interval of 95^th^ percentile (blue dots: Craig and Guillot, 2017 method; black asterisk: Bootstrap method). Only formulation types with minimum of 7 studies were included for this evaluation.

Differences of 95^th^ percentile between formulation groups and within each group using Bootstrap method to calculate lower and upper 95% confidence interval of the 95^th^ percentile for the dilutions are shown in **Figure 4**. Appendix C in Supplementary Material presents the 95^th^ percentile (95% upper confidence interval) using Bootstrap for each formulation type.

By analysing the overlap of the lower and upper 95% confidence interval of the 95^th^ percentile between groups, it’s observed that 95^th^ percentile of organic solvent formulation group is significantly different from water-based group for dilutions; however, solid formulation group is not significantly different from water-based and organic solvent groups. Within each group, EW shows the highest value among organic solvent formulations, SG the highest value among solid formulations, and CS the highest value among water-based formulations.

The difference in the width of the 95% confidence interval between the CS and ZC formulations between the two methods is likely related to several factors, such as sample size (n), data variability, and the use of the bootstrapping method:

a. sample size (n): sample size directly influences the precision of confidence interval estimates. In the Craig and Guillot, 2017 method, a smaller sample size may lead to wider confidence intervals reflecting greater uncertainty. In the case of bootstrapping, even with smaller sample sizes, it is possible to generate an empirical distribution of samples, thus providing more refined interval estimates, which may explain the narrower width for CS and ZC formulations;
b. data variability: the intrinsic variability of the dermal absorption measures can influence the observed differences between the methods. If CS and ZC formulations have considerable variability, the Craig and Guillot, 2017 method may tend to reflect this variability with wider confidence intervals. Bootstrapping, on the other hand, uses resampling, which can smooth out this variability, resulting in narrower confidence intervals;
c. use of bootstrapping: bootstrapping is a resampling-based technique that allows for the calculation of confidence intervals without assuming a specific data distribution. This can be particularly advantageous in contexts with high variability or non-parametric data. In the case of CS and ZC formulations, the bootstrap method can provide narrower intervals due to its ability to capture observed variability directly, without relying on parametric assumptions as the Craig and Guillot, 2017 method does.

In summary, bootstrapping offers a more flexible and potentially more accurate approach for estimates with smaller sample sizes and high variability which may explain the narrower confidence intervals for CS and ZC formulations.

### 4.4. Dermal absorption per formulation grouping

The 95% upper confidence interval of the 95^th^ percentile calculated with the two proposed methods (as described in 3.4.) are presented in Table 2.

**Table 2:** Dermal absorption 95^th^ percentile for different formulation groups.

| Tested form | Group | n<sup>(1)</sup> | 95<sup>th</sup> percentile<br>(95% upper confidence interval) |  |
| --- | --- | --- | --- | --- |
|  |  |  | Method presented by Craig and Guillot, 2017 | Bootstrap method |
| **Con cent rate** | Organic-solvent based | 2150 | 10.32 | 10.46 |
|  | Water-based | 2509 | 4.47 | 4.49 |
|  | Solid | 949 | 3.16 | 3.29 |
|  | Other types | 73 | 8.21 | 10.13 |
| **Dilu tion** | Organic-solvent based | 3318 | 41.73 | 41.98 |
|  | Water-based | 3463 | 36.58 | 36.60 |
|  | Solid | 1510 | 38.78 | 38.98 |
|  | Other types | 8 | NA | 3.98 |

(1) n: number of observations (individual values of replicate analysis).

The method presented by Craig and Guillot, 2017 might return ‘NA’ in situations where data is insufficient to obtain an upper confidence limit. This can also occur when the empirical distribution is highly skewed or when extreme quantiles are being estimated from small samples. To overcome this situation, the bootstrap method was used, which estimates the distribution by resampling with replacement from the original dataset (see Appendix B in Supplementary Material for details). This approach is useful when dealing with small samples or when the underlying distribution of the data is unknown or complex (DiCiccio and Efron, 1996, Justus *et al*., 2024).

### 4.5. Dilution factor effect on dermal absorption

This assessment was performed using the mean dermal absorption value for each dilution tested. Dilution factor between lower and higher dilutions were compared to the ratio of mean dermal absorption of these dilutions^7^. In total, 356 ratios of lower dilutions/higher dilutions are presented (“Dilution factor” *vs* “Fold of DA”) in **Figure 5**.

**Figure 5:**
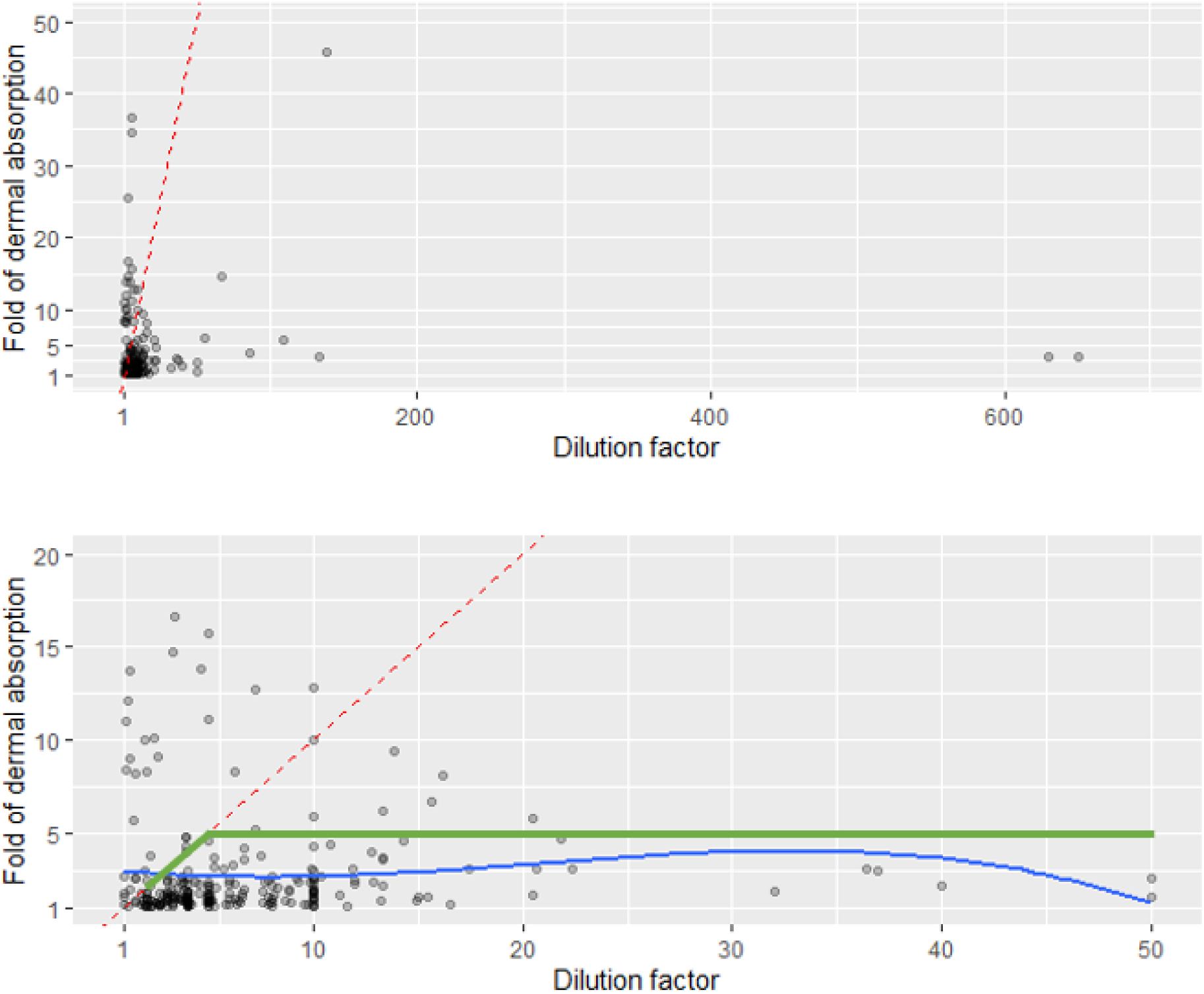
Dilution factor (x-axis) plotted against the fold of dermal absorption (y-axis) (only dilutions were considered) from combined ProHuma/ECPA dataset (top: whole range; bottom: zoom in on the region of interest). Blue line: polynomial regression from current dataset; Red line: regression according EFSA pro rata approach; Green line: regression according to the approach proposed by Aggarwal *et al.,* 2015. Intensity colour of points indicates higher numbers of observations.

No correlation between dilution factor and dermal absorption was found for the ProHuma/ECPA dataset, as represented by the blue line (polynomial regression) in **Figure 5**.

Pro rata correction applied by EFSA, 2017 for the extrapolation from a measured dilution to a higher non-measured dilution assumes a linear response and is represented as a red line. Aggarwal *et al*., 2015 proposed a linear correction between 2- and 5-fold dilutions and is represented as a green line.

Data from ProHuma/ECPA dataset demonstrates that the increase in dermal absorption with dilution is substantially less pronounced than predicted by the pro rata approach. It suggests a more nuanced pattern, where increases in dilution do not necessarily lead to proportional increases in dermal absorption. Notably, in 87% of cases, the increase in dermal absorption is less than 5-fold, even with significant dilution.

## 5. Discussions

The present assessment has confirmed previous findings (EFSA, 2017; Aggarwal *et al*., 2015) that organic-solvent formulations present higher relative dermal absorption values when compared to water-based and solid formulations, and that in general dilutions present higher dermal absorption values (in % of applied dose) when compared to concentrates.

### 5.1. Default values for dermal absorption

Based on the upper 95% confidence interval of 95^th^ percentiles obtained through bootstrapping – as described in Appendix B in Supplementary Material, proposed default dermal absorption values to support non-dietary risk assessment in Brazil are presented in Table 3. No default values nor extrapolation from another group are proposed to the ‘other type’ formulation group as the number of observations is extremely low and there’s a great variability on formulation types.

**Table 3:** Proposed default dermal absorption values when no experimental data is available.

| **Tested form** | **Group** | **Proposed default value (%) based on 95<sup>th</sup> percentile (95% upper confidence interval) using Bootstrap method** |
| --- | --- | --- |
| **Concentrate** | Organic-solvent based | 10 |
|  | Water-based | 4 |
|  | Solid | 3 |
| **Dilution** | Organic-solvent based | 42 |
|  | Water-based | 37 |
|  | Solid | 39 |

The main constraint of bootstrapping is the assumption that the sample is a good representation of the population. In the case of the combined ProHuma/ECPA dataset, the amount of data and its variability (in terms of active ingredients and formulation types) contributes to reduce bias in resampling. Additionally, bootstrapping is applied primarily to quantify uncertainty rather than as the main estimation tool, reducing over-reliance on its assumptions.

### 5.2. Extrapolation of experimental data

#### 5.2.1. Read-across between different formulations

When experimental data is available for a different formulation type than the one being assessed, two read-across possibilities are proposed, considering the statistical evaluation from the groupings and the dermal absorption differences within each group:

- read-across among different formulation group, using organic-solvent based dermal absorption values from experimental data as conservative surrogate for solid and water-based formulations;
- read-across within the same formulation group on a case-by-case basis, always using the highest upper 95% confidence interval of the 95^th^ percentile as the most conservative surrogate value to support such extrapolation (e.g., SE experimental dermal absorption data could be extrapolated to EC or OD as it presents higher upper 95% confidence interval of the 95th percentile, but it would not be justified to extrapolate SE experimental value to EW or ME.

These read-across proposals should be considered on a case-by-case basis, applying scientific judgement.

#### 5.2.2. Pro-rated correction between spray dilutions

According to the findings illustrated in **Figure 5**, the stepwise procedure for extrapolating dermal absorption data, as proposed by Aggarwal *et al*., 2015, is most suitable to the present dataset and it is detailed in Appendix D of the Supplementary Material. When experimental data for higher spray dilutions of a specific formulation type is unavailable, a pro-rated correction is applied if the evaluated dilution is between 2-fold and 5-fold more dilute than the highest tested dilution or until the corresponding default dermal absorption value (Table 3) is reached, whichever is lower.

## 6. Conclusions

A total of 468 studies from the ProHuma dataset were combined with the ECPA dataset, generating a ProHuma/ECPA combined dataset with 759 *in vitro* human skin dermal absorption studies, all compliant with GLP and the OECD 428 guideline. This resulted in a comprehensive and homogeneous database with a total of 14,803 observations, significantly increasing the number of observations per formulation type from previous works on the same subject. After evaluating this database using a cutting-edge machine learning method, including interpretable technique, it was concluded that it is sufficiently robust to support regulatory decisions on dermal absorption when specific studies are unavailable. This includes proposing default values and read-across possibilities in light of the implementation of the Brazilian regulation on non-dietary risk assessment.

## Credit authorship contribution statement

**Danilo Sarti:** conceptualization, statistical analysis including machine learning method, writing – reviewing; **Jörg Wagner:** data processing, initial statistical analysis, writing – reviewing; **Fabiana Palma:** conceptualization, project administration, writing – reviewing & editing; **Marcela Giachini:** writing – reviewing & editing; **Daniele Lautenschalaeger**: writing – reviewing & editing; **Janaina Pires**: writing – reviewing & editing; **Marina Sales**: writing – reviewing & editing; **Patricia Faria**: writing – reviewing & editing; **Vanessa Lupianhez**: writing – reviewing & editing; **Heloisa Kalvan**: conceptualization, writing – reviewing & editing; **Marine Le Bras:** data processing, initial statistical analysis; **Lisa Bertomeu:** data processing, initial statistical analysis.

## Acknowledgement

ProHuma is grateful to all the company members that supported this project, to the company members that contributed by providing data from dermal absorption studies to compose the dataset analyzed in this project, and to the authors – scientific specialists of the ProHuma technical committee.

ProHuma also would like to thank to the Toxicology General Office – GGTOX from ANVISA for the valuable support through recommendations for improvements throughout the development of this project.

## Supplementary Material

The following is the Supplementary data to this project.

### Appendix A. Data collection (Top) and steps involved in the data processing (Bottom)

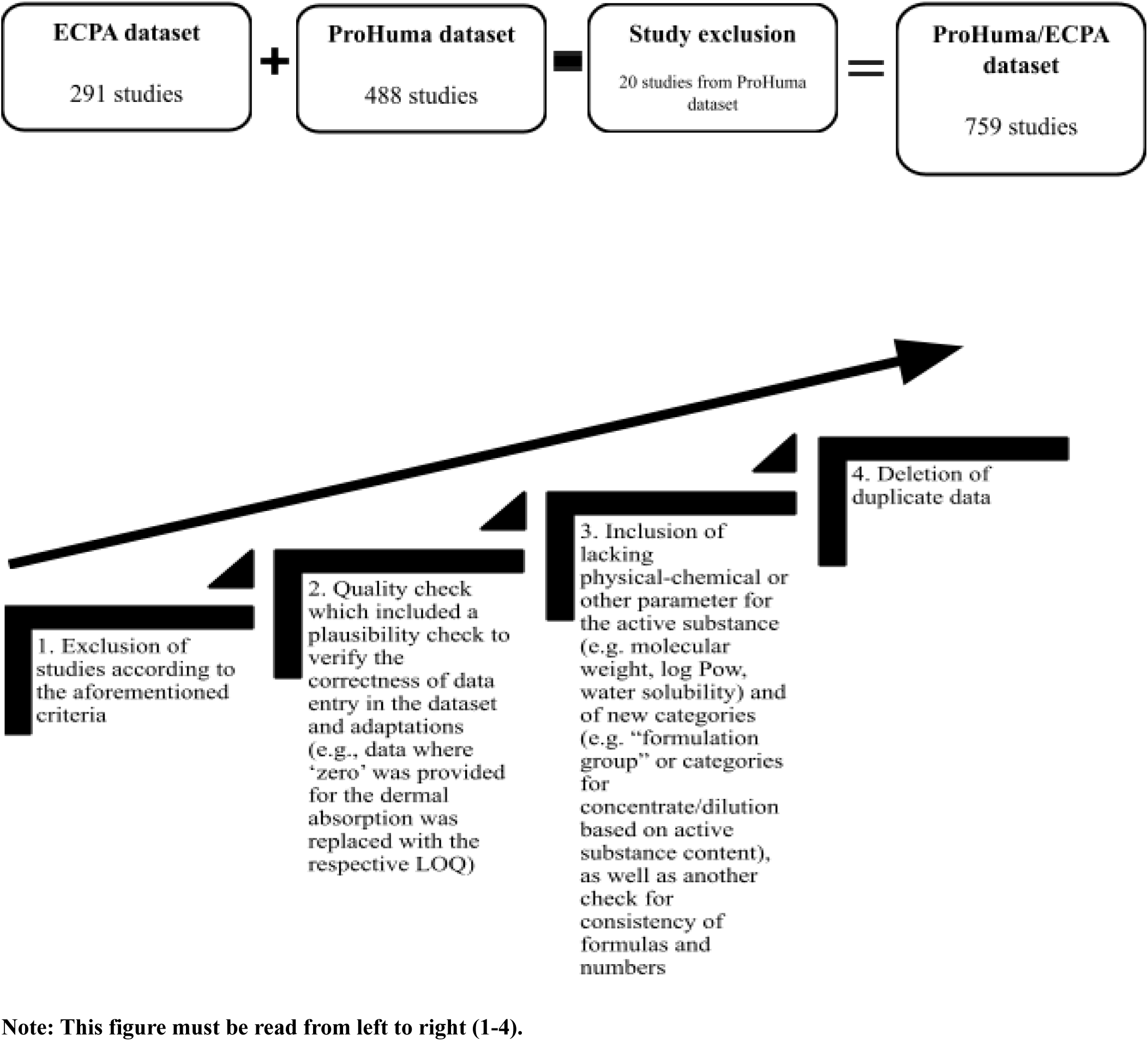

### Appendix B. Description of the method implemented by R function

Given a sample x = {x_1, x_2, …, x_n} of size n, the goal is to estimate the quantile q_alpha and the corresponding confidence interval. The function operates as follows:

1. **Calculate the empirical quantile**: For a given quantile level alpha, the empirical quantile is the value x_r such that P(q_alpha <= x_r) <= beta, where beta is the desired confidence level;
2. **Determine the rank r**: The rank r is chosen such that the probability of the quantile being less than or equal to the r-th order statistic is greater than or equal to beta.

The function ‘ub.leftCI’ as presented in Craig and Guillot, 2017 might return ‘NA’, as describe in Table B7 from EFSA, 2017 in situations where the sample size is too small, particularly when the confidence level beta is very high, and the required rank r exceeds the sample size. This can occur when the empirical distribution is highly skewed or when extreme quantiles (e.g., 99^th^ percentile) are being estimated from small samples. To copy with such situation, it was also used an alternative approach of estimating the upper bound of a confidence interval using a bootstrap method (DiCiccio and Efron, 1996, Etzioni *et al*., 2020, Justus *et al*., 2024).

The bootstrap method offers an alternative approach to estimating confidence intervals, particularly when the assumptions of the ‘ub.leftCI’ function are not met or when the sample size is inadequate.

The bootstrap procedure involves the following steps:

1. **Resampling**: From the original sample x, generate a large number B of bootstrap samples x^*_b (for b = 1, 2, …, B) by resampling with replacement;
2. **Estimate the quantile for each sample**: For each bootstrap sample, compute the desired quantile q_alpha.
3. **Construct the confidence interval**: The confidence interval is constructed by taking the empirical quantiles from the distribution of the bootstrap estimates {q*_{alpha,1}, q*_{alpha,2}, …, q{alpha,B}}.

In the present analysis, it was implemented a non-parametric bootstrap method with 100,000 resamples, drawing samples with replacement from the observed dataset. For each resample, it was computed the 95^th^ percentile (q95) and other relevant quantiles. The bootstrap confidence intervals were then obtained using the percentile method, where the 2.5^th^ and 97.5^th^ percentiles of the bootstrap distribution served as the bounds for the 95% confidence interval.

### Appendix C. Effect of formulation type

**Table 1:** Overview of dermal absorption’s percentiles (%) by formulation types for concentrates and dilutions.

| Formulation type | Number of observations |  | 95<sup>th</sup> percentile (95% upper confidence interval) – Bootstrap method |  |
| --- | --- | --- | --- | --- |
|  | Concentrates | Dilutions | Concentrates | Dilutions |
| **Organic solvent** |  |  |  |  |
| **EC** | 1413 | 2146 | 12.30 | 41.85 |
| **EW** | 144 | 224 | 20.63 | 56.67 |
| **ME** | 74 | 135 | 6.48 | 52.89 |
| **OD** | 157 | 268 | 11.39 | 33.16 |
| **SE** | 362 | 545 | 6.67 | 47.69 |
| **Water-based/dispersed** |  |  |  |  |
| **CS** | 72 | 102 | 7.33 | 56.37 |
| **FS** | 474 | 311 | 3.30 | 16.09 |
| **SC** | 1651 | 2566 | 3.67 | 39.54 |
| **SL** | 251 | 416 | 9.99 | 32.48 |
| **ZC** | 61 | 68 | 7.65 | 38.16 |
| **Solid** |  |  |  |  |
| **SG** | 60 | 66 | 5.18 | 71.67 |
| **WG** | 816 | 1338 | 3.06 | 36.96 |
| **WP** | 73 | 106 | 6.98 | 41.29 |

### Appendix D. Procedure for extrapolation of dermal absorption data (reproduced from Aggarwal *et al*., 2015)

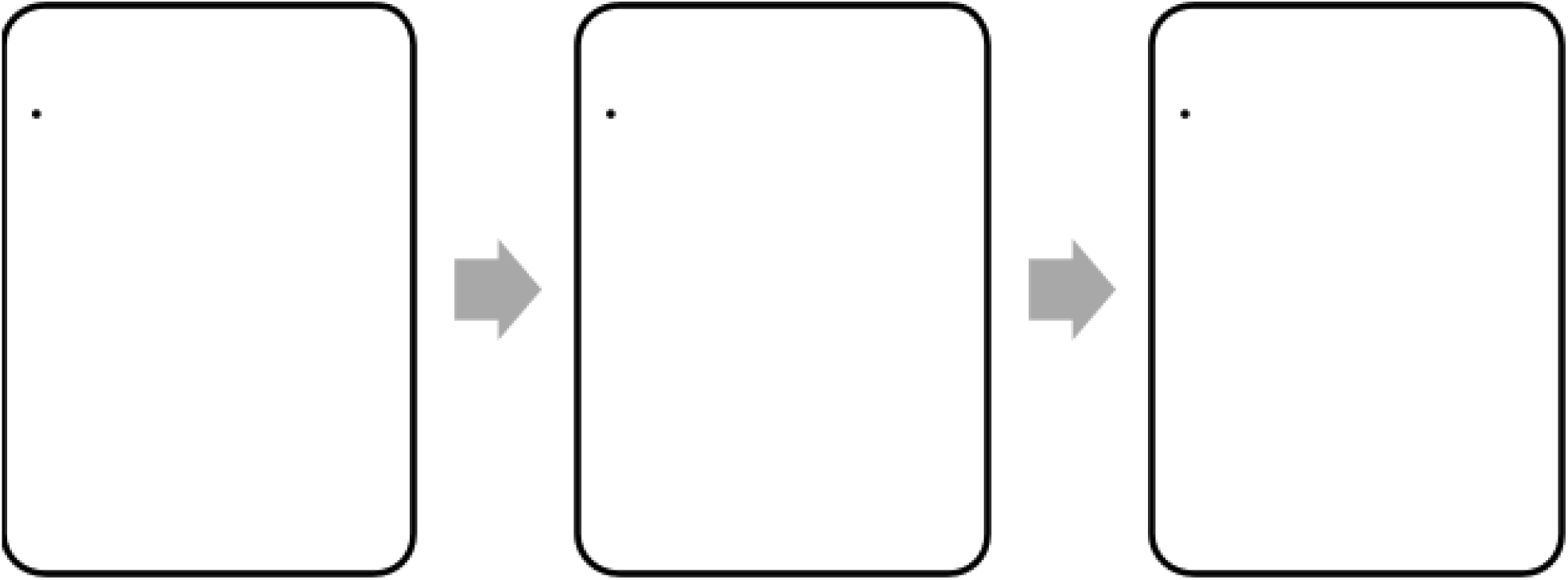

## Footnotes

1 Throughout the text, the terms ‘pesticide’, ‘agrochemical’, and ‘plant protection product’ (PPP) will be used interchangeably.

2 Formulation types from this grouping as per EFSA, 2017: emulsifiable concentrate (EC), emulsion, oil in water (EW), suspo-emulsion (SE), dispersible concentrate (DC), oil miscible liquids (OL/OF), oil-based suspension concentrates (OD), emulsion for seed treatment (ES), microemulsion (ME).

3 Bait concentrate (CB), capsule suspension (CS), gel for direct application (GEL/GD), bait, ready for use (RB), mixture of capsule suspension and suspension concentrate (ZC), seed coated with a pesticide (PS), experimental solution of active substances in solvent (AI).

4 Soluble concentrate (SL), suspension concentrate (SC), flowable concentrate for seed treatment (FS), flowable (FL) (=SC).

5 Wettable powder (WP), water-dispersible granules (WG/WDG), water-soluble granules (SG), water-soluble powder (SP), powder for dry seed treatment (DS).

6 In this context, an observation means a reported dermal absorption individual result of each replicate analysis.

7 Dilution factor = concentration [g/L] of lower dilutions / concentration [g/L] of higher dilution; Fold of DA = DA of dilution / DA of concentrate

